# Barley powdery mildew effector CSEP0162 targets multivesicular body-associated MON1 important for immunity

**DOI:** 10.1101/2022.05.06.490921

**Authors:** Wenlin Liao, Mads E. Nielsen, Carsten Pedersen, Wenjun Xie, Hans Thordal-Christensen

## Abstract

Encasements formed around haustoria and biotrophic hyphae as well as hypersensitive reaction (HR) cell death are essential plant immune responses to filamentous pathogens. Here we study a possible reason why these responses are absent in susceptible barley attacked by the powdery mildew fungus. We find that the effector CSEP0162 from this pathogen targets plant MON1, important for fusion of multivesicular bodies to their target membranes. Over-expression of CSEP0162 and silencing of barley *MON1* both inhibit encasement formation. We find that the Arabidopsis ecotype No-0 has partial resistance to powdery mildew, and that this is dependent on MON1. Surprisingly, we find the MON1-dependent resistance in No-0 not only include an effective encasement response, but also HR. Similarly, silencing of MON1 in barley also blocked *Mla3*-mediated HR-based powdery mildew resistance. These data indicate that MON1 is a vital plant immunity component, and we speculate that the barley powdery mildew fungus introduces the effector CSEP0162 to target MON1 and reduce encasement formation and HR.

**Highlight:** MON1 is essential for MVB fusion to plasma membrane. We find that MON1 also is important for immunity, and that it is targeted by the barley powdery mildew effector CSEP0162.

## Introduction

The plant immune system is activated in individual steps during the process of pathogen attack. Initially, plant plasma membrane (PM) receptor kinases detect pathogen-associated molecular patterns and subsequently activate pattern-triggered immunity (PTI) (Zipfel and Oldroyd, 2017). As a countermeasure, the pathogens introduce effector molecules into the plant cell cytosol to prevent activation of PTI. However, these effectors may be recognized either directly or indirectly by plant nucleotide-binding leucine-rich repeat (NLR) receptors, whereby effector-triggered immunity (ETI) is activated resulting in a programmed cell death (Jones *et al*., 2016; Thordal-Christensen, 2020; Kanja and Hammond-Kosack, 2020; Ngou *et al*., 2022). PTI and ETI responses consist partly of a complex transcriptional reprogramming and partly of cellular responses. The latter include papillary cell wall appositions at sites of attack, encasements in the form of cell wall extensions that enclose pathogen structures invading the plant cell, and the hypersensitive reaction (HR) programmed cell death. Papillae and encasements, block penetration into the plant cell and nutrient transfer from the pathogen, respectively, while the HR is detrimental to biotrophic pathogens depending on living plant cells.

From previous studies, several Arabidopsis proteins have been found important for penetration resistance towards the barley powdery mildew fungus (*Blumeria graminis* f.sp. *hordei*, *Bgh*), including the so-called PEN proteins (Hématy *et al*., 2020). The syntaxin PEN1 (SYP121), as well as its barley orthologue ROR2, are required for timely papilla formation (Assaad *et al*., 2004; Böhlenius *et al*., 2010). Meanwhile, PEN1 and its closest homologue, SYP122, have a shared function in papilla and encasement formation *per se* (Rubiato *et al*., 2022). These syntaxins are likely required for fusion of multivesicular bodies (MVBs) to the PM at the site of fungal attack to mediate papilla and encasement formation. Both these structures contain extracellular vesicles (EVs) secreted when MVBs fuse with the PM. These EVs are labelled with PEN1 (Nielsen *et al*., 2012; 2017; Rutter and Innes, 2017) probably because this syntaxin is carried onto the intraluminal vesicles (ILVs) as they form in the MVBs. Meanwhile, only EV secretion into encasements, and not into papillae, is dependent on VPS9a, a Rab guanine-nucleotide exchange factor (GEF) required for activation of the Rab5 GTPases (Nielsen *et al*., 2017). GTP-bound Rab5 GTPases are known to regulate the maturation of ESCRT-dependent MVBs and furthermore recruit the MON1/CCZ1 heterodimer that serves as a GEF activating Rab7 GTPases (Cui et al., 2014). Completion of the Rab5➔Rab7 transition is essential for the fusion of MVBs with the target membrane being the tonoplast, surrounding the vacuole, or alternatively the PM. Consequently, VPS9a is predicted to have a role in encasement formation as it via activation of Rab5 recruits MON1/CCZ1 to activate Rab7, which is essential for fusion of MVBs to the PM (Hansen and Nielsen, 2018). In Arabidopsis Col-0, *MON1* knock-out (KO) mutants, *mon1-1* (=*sand-1*) and *sand-2*, were found to suffer from very poor germination and impaired growth, whereas Nossen-0 (No-0) KO *mon1-2* plants are intermediate in size (Cui *et al*., 2014; Ebine *et al*., 2014; Singh *et al*., 2014), confirming the fundamental importance of this gene. Interestingly, Ortmannová *et al*. (2022) recently studied Arabidopsis EXO70 complexes important for vesicle tethering to the PM. Here, they identified the EXO70B2 complex to interact with PEN1 and to be required for normal papilla and encasement formation. Moreover, they also uncovered an interaction between EXO70B2 and RabG3C, an Arabidopsis Rab7 homologue, suggesting that PEN1 mediates an MVB-PM fusion regulated by the EXO70B2 complex during papilla and encasement formation.

Powdery mildew fungi are serious pathogens on numerous plant species, and their autonomous attack on individual leaf epidermal cells makes these plant-pathogen interactions useful for cellular studies. Powdery mildew fungi are biotrophic pathogens that take up nutrients from the plant by help of haustoria inside the host cells. The genome of *Bgh* encodes hundreds of candidate secreted effector proteins (CSEPs) to promote the attack (Pedersen *et al*., 2012; Frantzeskakis *et al*., 2018). Yet, very few of these have been studied for their contribution to fungal virulence, let alone their plant target proteins. Here, it is expected that some CSEPs hamper papilla and encasement formation in the barley host plant. In Arabidopsis, haustoria of the non-adapted *Bgh* are generally encased (Nielsen *et al*., 2017). In contrast, barley does not encase *Bgh* haustoria, despite the cellular machinery for making encasements exists. This appears from the facts that this structure can be stimulated after treatment with the ergosterol biosynthesis inhibiting fungicide, tetraconazole (Maffi *et al*., 1995; Bolton *et al*., 2016). Therefore, we assume *Bgh* secretes CSEPs to effectively inhibit encasement formation.

CSEP0162 was previously found to interact with small heat shock proteins (sHSPs) (Ahmed *et al*., 2015). In the present work, we re-screened for more barley target proteins of CSEP0162 using yeast 2-hybrid (Y2H) and found that it also targets MON1. Over-expression of CSEP0162, as well as silencing of *MON1*, hampered encasement formation. Similar results were found in the loss-of-function mutant *mon1-2* of Arabidopsis, showing that MON1 serves a conserved function in encasement formation. More surprisingly, silencing of *MON1* was found to hamper barley NLR-mediated powdery mildew resistance. Similarly, *mon1-2* in Arabidopsis also hampered a cell death reaction induced by the powdery mildew fungus, suggesting that HR can be at least partially dependent on MON1. In support of MON1 being a likely effector target also in Arabidopsis, we provide evidence that the Col-0 *mon1-1* lethality is partly due to EDS1-dependent autoimmunity.

## Materials and Methods

### Plant material

The following barley (*Hordeum vulgare*) lines were used in this work. *ror2*, a syntaxin mutant in cv. Ingrid w. low penetration resistance (Collins *et al*., 2003). The near-isogenic cv. Pallas lines, P-01 and P-02, with the powdery resistance genes, *Mla1* and *Mla3*, respectively (Kølster *et al*., 1986). Barley plants were grown under 16 h (20°C) day (150 μE m^-2^ s^-1^), 8 h night (15°C). The following *Arabidopsis thaliana* lines were used in this work. Ecotype Columbia-0 (Col-0) and its mutants, *mon1-1* (T-DNA insertion line SALK_075382, Singh *et al*., 2015), *eds1-2* (Bartsch *et al*., 2006), *ndr1-1* (Century *et al*., 1997). Ecotype Nossen-0 (No-0) and its mutant *mon1-2* (Ds transposon line 54-4894-1 received from RIKEN) (Cui *et al*., 2014). Arabidopsis plants were grown under 8 h (21°C) day (125 μE m^-2^ s^-1^), 16 h night (15°C). Arabidopsis germination rates were determined on ½ MS phytoagar. Mutant allele genotypes were determined by PCR using primers listed in Supplementary Table S1.

### Fungal material

The barley powdery mildew fungus (*Bgh*) isolates, A6 and C15, were propagated on P-01 and P-02, respectively, by weekly transfer. The Arabidopsis powdery mildew fungus, *Golovinomyces orontii* (*Go*) isolate MPIPZ was propagated on *eds1-2* by bi-weekly transfer.

### Construction of plasmids

Coding sequences of *HvMON1*, *HvMON1i* and *CSEP0162* were amplified from *Bgh-* inoculated barley (Ingrid) cDNA by Q5^®^ High-Fidelity DNA Polymerase (NEB). The primers are listed in Supplementary Table S1. The PCR products were cloned into pDONR221 by BP Clonase (Invitrogen). Inserts were recombined into destination vectors listed in Supplementary Table S2 by Gateway LR clonase (Invitrogen) reactions. All constructs were confirmed by sequencing.

### Yeast Two-Hybrid Screen

A barley cDNA library, made from *Bgh*-infected barley leaves and cloned into vector pDEST-ACT2 (Zhang *et al*., 2012), was screened using the *Bgh* effector pDEST-AS2-CSEP0162 bait-construct. The yeast, *Saccharomyces cerevisiae*, transformation protocol and recipe of synthetic dropout (SD) media, described by Zhang *et al*. (2012), was used to transform the prey library and the bait construct into yeast strains Y8800 and Y8930, respectively. These two haploid strains have different mating types and they have the mutations *ade2*, *his3*, *leu2* and *trp1*. The strains were mated by mixing 1 ml aliquot of the library strain and 5 ml of the bait strain into 45 ml 2×YPDA medium in a 2 L flask and incubated at 30°C overnight at 30-50 rpm. The overnight culture was plated onto SD medium without tryptophan, leucine, histidine, and adenine with 2.5 mM 3-AT. After 2 days of incubation at 30°C, the largest yeast colonies were picked for colony PCR and sequencing of the prey insert. From yeast colonies with prey constructs encoding barley proteins of more than 20 amino acids in frame with the Gal4 activation domain, the plasmids were extracted and retransformed into Y8800 to confirm the interaction. SNF1 (Y8930) and SNF4 (Y8800) were used as a positive control (Durfee et al., 1993), and negative controls were pDEST-AS2-CSEP0105 (Ahmed *et al*., 2015) and empty vector.

### Bimolecular fluorescence complementation (BiFC) and protoplast protein co-localization

The set of Gateway binary Ti destination vectors with nGFP or cCFP and different fusion orientations, generated by Kamigaki *et al*. (2016), were used. Full-length coding sequences of *Hv*MON1 and CSEP0162 (without signal peptide) were cloned into the BiFC vectors by LR reactions and confirmed constructs were introduced into *Agrobacterium tumefaciens* strain GV3101. Overnight cultures were harvested and resuspended to OD_600_=0.7 in 10 mM MgCl_2_, 10 mM MES, 0.1 mM Acetosyringone. A total of eight combinations of *A. tumefaciens* with constructs for *Hv*MON1 and CSEP0162 fused to complementary GFP-fragments in different orientations were co-infiltrated into *Nicotiana benthamiana* leaves. Dimerization of 14-3-3 was used as positive control (Aitken, 2006).

Protoplasts were isolated from the second true leaves of 7-day-old barley and transformed according to Saur *et al*. (2019).

GFP signal in *N. benthamiana* 2 days after infiltration, and mYFP and mCherry signals in protoplasts after overnight incubation in darkness were detected by Leica SP5 confocal microscopy (GFP, excitation: 488 nm, emission: 518-535 nm; mYFP, excitation: 513 nm, emission: 526-555 nm; mCherry, excitation: 588 nm, emission: 613-650 nm) at the Centre for Advanced Bioimaging, University of Copenhagen.

### Barley epidermal cell transient induced gene silencing (TIGS) and over-expression

*Hv*MON1 was transiently induced gene silenced (TIGS) as described by (Douchkov *et al*., 2005). An *Hv*MON1 RNAi fragment (316 bp) was designed by the siRNA-Finder (si-Fi) Software (Lück et al., 2019) and introduced twice in opposing orientation in the destination vector pIPKTA30N to produce a hairpin transcript (Douchkov *et al*., 2005). A construct for over-expression of YFP-CSEP0162 was previously generated by Ahmed et al. (2015). Each of these pIPKTA30N-*HvMON1i* and pUbi-YFP-CSEP0162-nos constructs were cotransformed with pUbi-GUS-nos as marker into barley epidermal cells by particle bombardment according to Smigielski *et al*. (2019). Subsequently, the leaves were placed in closed 1% phytoagar plates w. 10 μg ml^-1^ benzimidazole at 16 h (20°C) day (150 μE m^-2^ s^-1^), 8 h night (15°C). To study transformed cells, leaves were stained with X-Gluc for GUS activity according to Douchkov *et al*. (2005),

### Immune response scorings

To induce encasements around *Bgh* haustoria, barley leaves were sprayed with 100 μg ml^-1^ tetraconazole in 20% acetone, 0.04% Tween-20 (Maffi *et al*., 1995) 2 hours before inoculation with *Bgh*. To study encasements, either alone or in combination with GUS staining, they were visualized 5 days after inoculation by their callose content after staining of the leaves with 0.01% aniline blue in 1 M glycine, pH 9.5, and UV epifluorescence microscopy.

Penetration rate, encasement formation, HR cells and fungal development in Arabidopsis were scored by light and UV epifluorescence microscopy as described by Nielsen *et al*. (2017). In short, for scoring of penetration success, leaf material was trypan blue stained at 2 days after *Bgh* or *Go* inoculation. For each leaf, a minimum of 50 penetration attempts (presence of a fungal appressorium) were scored using light microscopy. Penetration was determined by the presence of a fungal haustorium. Callose staining was performed as above.

### RNA Extraction and Quantitative RT-PCR

Total RNA was extracted using Monarch^®^ RNA Cleanup Kit (NEB). Reverse transcription and cDNA synthesis were performed using NEBNext^®^ RNA First Strand Synthesis Module (NEB). Transcript quantification was carried out using Stratagene MX3000P real-time PCR detection system (Agilent Technologies) with FIREPol^®^ EvaGreen^®^ Mix (Solis BioDyne). Primers used to amplify PCR products of maximum 200 bp are described in Supplemental Table S1. The ubiquitin conjugating factor (UBC2) was used as barley reference gene (Skov *et al*., 2007). The level of gene expression was calculated using the relative quantification (^ΔΔ^Ct) algorithm by Livak and Schmittgen (2001) based on two technical and three biological replicates.

### Barley Stripe Virus Induced Gene Silencing (VIGS)

The tripartite genome of Barley Stripe Mosaic Virus (BSMV) was used as basis for barley gene silencing. The binary Ti-constructs pCaBS-α, pCaBS-β, pCa-γbLIC and pCaBS-γb-*TaPDS* were described by Yuan et al. (2011). The Gateway cassette was inserted into the ligation-independent cloning site of pCa-γbLIC, and an RNAi sequences of *HvMON1* and a full-length coding sequence of *mYFP* were inserted by Gateway LR clonase (Invitrogen) reactions. All constructs were transformed into *A. tumefaciens* strain EHA105 by selection on rifampicin (25/μg ml) and kanamycin (100 μg/ml). Confirmed strains were co-infiltrated into *N. benthamiana* with *A. tumefaciens* containing pCaBS-α and pCaBS-β according to the method described above. When BSMV symptom appeared on upper leaves approximately 10 dpi, then BSMV infected leaves were collected and ground in 20 mM Na-phosphate, pH 7.2 with 1% silica. The homogenates were smeared onto first leaves of 7-day-old barley seedlings by rubbing gently with fingers. The third leaves of treated barley plants were collected about two weeks later for qRT-PCR or *Bgh* inoculation. Silencing of phytoene desaturase using pCaBS-γb-*TaPDS* was used as a positive indicator for the VIGS system, while pCa-γb-mYFP was used as negative control.

## Results

### CSEP0162 interacts with barley MON1

The *Bgh* effector candidate, CSEP0162, was previously found to contribute to fungal virulence and to interact with sHSPs (Ahmed *et al*., 2015). In a search for additional barley target proteins of CSEP0162, the Y2H library used in Ahmed et al. (2015) was re-screened, leading to identification of a prey clone encoding the C-terminus of *Hv*MON1 (amino acids 518-577). In support of this, the full-length *Hv*MON1 was found also to interact with CSEP0162 in Y2H (Fig. 1A). Interaction between these two proteins occurred *in planta* as well shown by BiFC following agroinfiltration of *Nicotiana benthamiana* leaves. Here the nGFP-*Hv*MON1/CSEP0162-cGFP combination reconstituted a fluorescent protein (Fig. 1B), whereas the other seven combinations did not.

**Figure 1.**
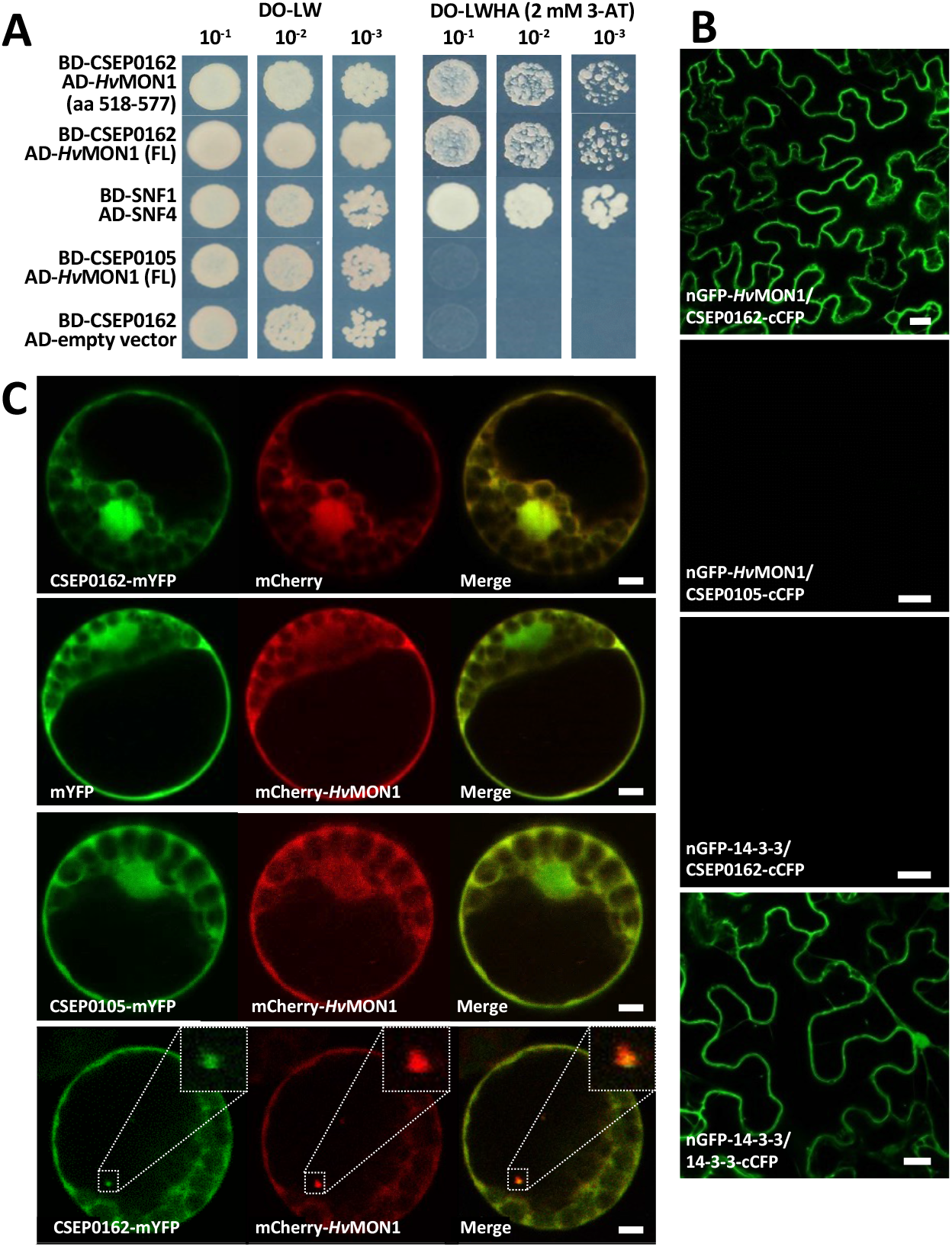
CSEP0162 interacts with HvMON1. **(A)** Yeast two-hybrid (Y2H). Yeast transformed with constructs fusing CSEP0162 with the GAL4 <u>b</u>inding-<u>d</u>omain (BD) and full-length (FL) *Hv*MON1 with the <u>a</u>ctivation-<u>d</u>omain (AD). Growth on <u>d</u>rop<u>o</u>ut (DO)-medium lacking leucine (L) and tryptophan (W) indicated presence of both constructs. Growth on DO-medium lacking L, W, histidine (H) and adenine (A) w. 2 mM <u>3</u>-<u>a</u>mino-1,2,4-<u>t</u>riazole (3-AT) indicated protein-protein interaction. SNF1/SNF4 was used as a positive control. CSEP0105 and empty vector were used as negative controls. **(B)** Bimolecular fluorescence complementation (BiFC) in *N. benthamiana* leaves after *Agrobacterium* infiltration, and epidermal cells were observed by laser scanning confocal microscopy (LSCM). The GFP signal was captured 48 h after co-expressing nGFP-HvMON1 and CSEP0162-cCFP. CSEP0105 and 14-3-3 were used as negative controls. Dimerization of 14-3-3 protein was used as a positive control. Size bars, 20 μm. **(C)** Co-expression of CSEP0162-mYFP and mCherry-*Hv*MON1 in barley P-02 protoplasts observed by LSCM 24 h after transformation. Size bars, 5 μm.

To study the interaction further, GFP-CSEP0162 and mCherry-*Hv*MON1 were coexpressed in barley mesophyll protoplasts. As reported by Ahmed *et al*. (2015), GFP-CSEP0162 localized in the nucleus and cytoplasm, whereas mCherry-*Hv*MON1 was only visible in the cytoplasm (Fig. 1C). Strikingly, CSEP0162 and *Hv*MON1 co-localized in diffuse ~1 μm structures (inserts in Fig. 1C, Supplementary Video S1). Similar structures were never observed when CSEP0162 and *Hv*MON1 were expressed individually. Ahmed et al. (2015) observed somewhat larger diffuse structures when co-expressing CSEP0162 and interacting sHSPs and referred to these as aggresomes. Indeed, sHSPs have the ability to enter aggresome formation together with interacting proteins (Johnston and Samant, 2021; Reinle *et al*., 2022). We suggest the CSEP0162/*Hv*MON1 structures also to be aggresomes based on their diffuse nature, and we take this co-localization as additional evidence for molecular interaction between these proteins.

### Hv*MON1 and CSEP0162 regulate encasement formation around powdery mildew haustoria in barley*

Previously, we have shown that VPS9a is required for the correct formation of encasements around powdery mildew haustoria in Arabidopsis (Nielsen *et al*., 2017). While MON1 acts downstream of VPS9a to activate Rab7 to mediate MVB fusions to the tonoplast (Cui *et al*., 2014; Ebine *et al*., 2014; Singh *et al*., 2014), we envisage that this pathway also mediates MVB to PM fusion. Therefore, we used RNA interference (RNAi)-based transient-induced gene silencing (TIGS) to test whether *Hv*MON1 is required for encasement formation around *Bgh* haustoria in barley. Here we made use of tetraconazole to stimulate encasements in the barley epidermal cells (Supplementary Fig. S1) and found that TIGS of *Hv*MON1 reduced the formation of this defense structure by more than 70% (Fig. 2A-B). Since CSEP0162 interacts with *Hv*MON1, we speculated that this effector would influence encasement formation as well. Therefore, we over-expressed CSEP0162 in the same set-up and found a 50% reduction in encasement formation (Fig. 2C). In summary, our data suggest that CSEP0162 targets *Hv*MON1 to inhibit encasement formation.

**Figure 2.**
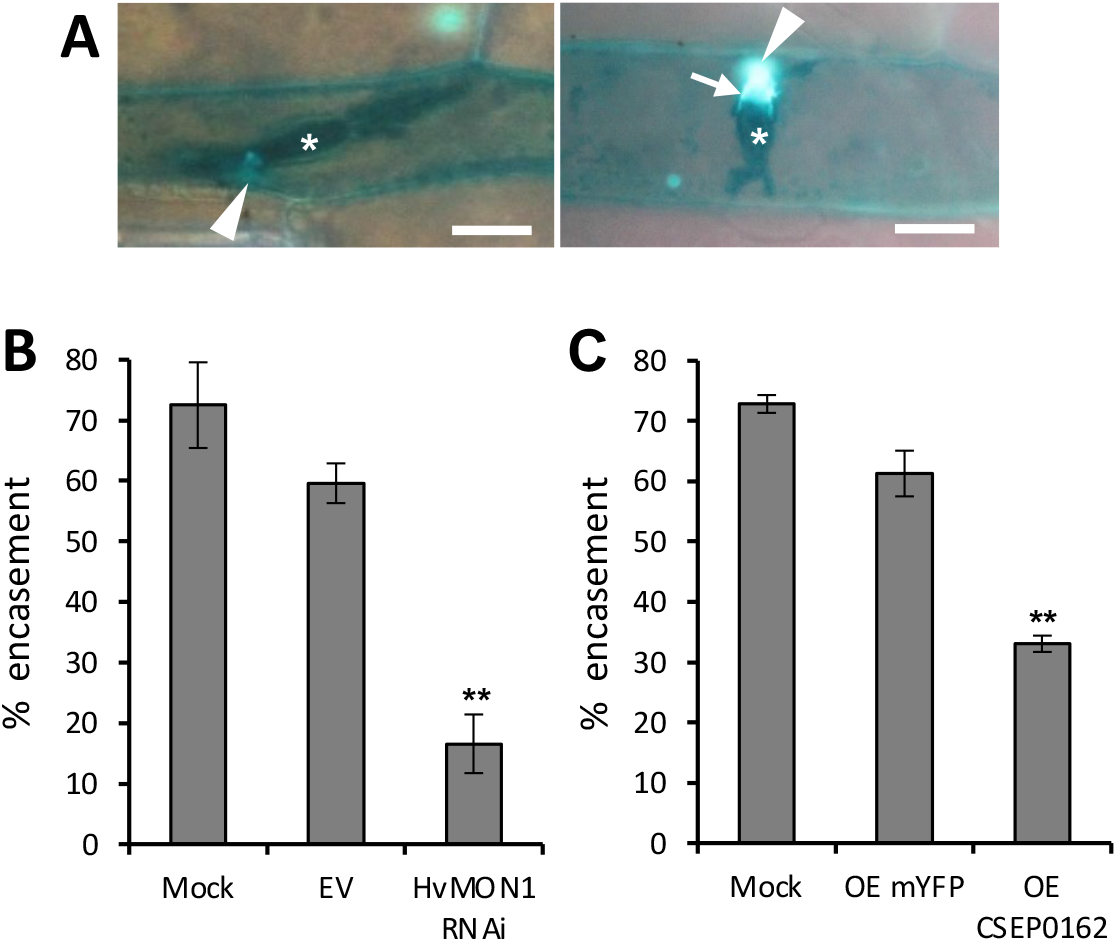
Encasement of *Bgh* haustoria requires *Hv*MON1 and is suppressed by CSEP0162. **(A)** *Bgh* (C15) haustoria (aster) in barley *ror2* epidermal cell without (left) and with callose-containing encasement (arrow) (right) imaged by UV-fluorescence microscopy after aniline blue treatment. Size bars, 20 μm. **(B,C)** Effect of *MON1* silencing and CSEP0162 over-expression (OE) on encasement of *Bgh* (C15) haustoria. Mock: data from untransformed cells of leaves with *MON1* RNAi and OE CSEP0162 cells, respectively. **(A-C)** Leaves of eight-day-old barley plants were transformed by particle bombardment. Two days later, they were treated with tetraconazole and *Bgh* inoculation. Encasement scorings were made after another 5 days. The data are mean values of four experiments. Error bars, SE. **, *P*<0.01 assessed by Student’s T-tests. n=4.

### Arabidopsis MON1 is required for immunity

MON1 is part of an evolutionarily conserved transport system that enables fusion of mature MVBs to their target membrane (Cui *et al*., 2014; Ebine *et al*., 2014; Singh *et al*., 2014). In support of this, we found that over-expressing GFP-*Hv*MON1 in the Arabidopsis Col-0 *mon1-1* KO mutant reverts this line’s phenotype to normal (Supplementary Fig. S2A-B). In roots, the GFP-*Hv*MON1 signal displayed a distinct punctate pattern, which in response to the PI3-kinase inhibitor wortmannin became ring-like, consistent with previous observations that *At*MON1 localizes to the MVB (Supplementary Fig. S2C; Singh *et al*., 2014). Having confirmed that *Hv*MON1 is the functional orthologue of Arabidopsis MON1, inspired us to study whether MON1 has the same role in immunity in Arabidopsis as what we observed in barley. For this purpose, we turned to use the ecotype No-0. In this line, the *mon1-2* KO mutant has superior growth relative to the Col-0 *mon1-1* mutant, including that homozygous *mon1-2* can grow to maturity and seed set (Cui *et al*., 2014). Moreover, while seed germination is largely impaired in *mon1-1* (Col-0) (see below), the overall seed germination rate was unaffected in *mon1-2* (No-0). Like *Bgh* on barley, *Go* is an example of a pathogen that has adapted to overcome the immunity presented by Arabidopsis, even in wild-type plants (Spanu *et al*., 2010). However, in comparison to Col-0, we discovered that No-0 is partially resistant to the *Go* powdery mildew fungal isolate, MPIPZ (Fig. 3A). Strikingly, this resistance was severely compromised in *mon1-2* (Fig. 3A-B), showing that MON1 is essential for immunity also in Arabidopsis.

**Figure 3.**
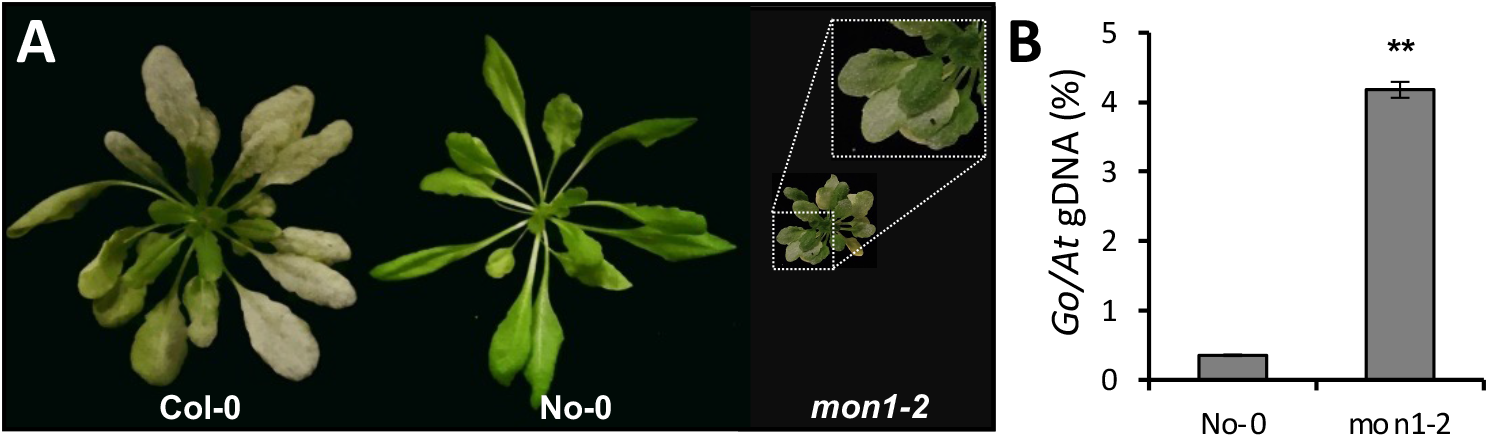
Resistance of No-0 *to Go* requires *At*MON1. **(A)** No-0, in contrast to Col-0, has no visible powdery mildew symptoms 7 days after inoculation with *Go*. This resistance of No-0 was broken by the *mon1-2* mutation. **(B)** qPCR-based quantification of fungal biomass 4 dpi. Error bars, SE. **, *P*<0.01 assessed by Student’s T-tests. n=3.

### MON1 is required for normal penetration resistance, encasement formation and for HR in Arabidopsis

To study the role of MON1 in the resistance displayed by No-0 further, we analyzed the initial stages of powdery mildew attack microscopically. Here we found that *Go* quite efficiently succeeds in penetrating No-0, nonetheless, the penetration rate in *mon1-2* was marginally increased (Fig. 4A). Furthermore, we found that the average length of secondary hyphae from successfully penetrating spores was longer on leaves of *mon1-2* than No-0, indicating that post-invasive immunity of *mon1-2* is hampered (Fig. 4B). In support of this, we found a reduction in the encasement response by more than 80% (Fig. 4C-D). The lack of encasements not only explains the increase of the growth rates of secondary hyphae in *mon1-2*, but also shows that encasement formation in monocots (Fig. 2B) and dicots has a common requirement for MON1. In addition, we found that more than 30% of the successfully penetrated host cells in No-0 had initiated an HR. Remarkably, this was reduced by more than 90% in *mon1-2* (Fig. 4E-F). Combined, this shows that in the No-0 ecotype, MON1 is crucial for resistance based on the immune responses of encasement formation and HR cell death.

**Figure 4.**
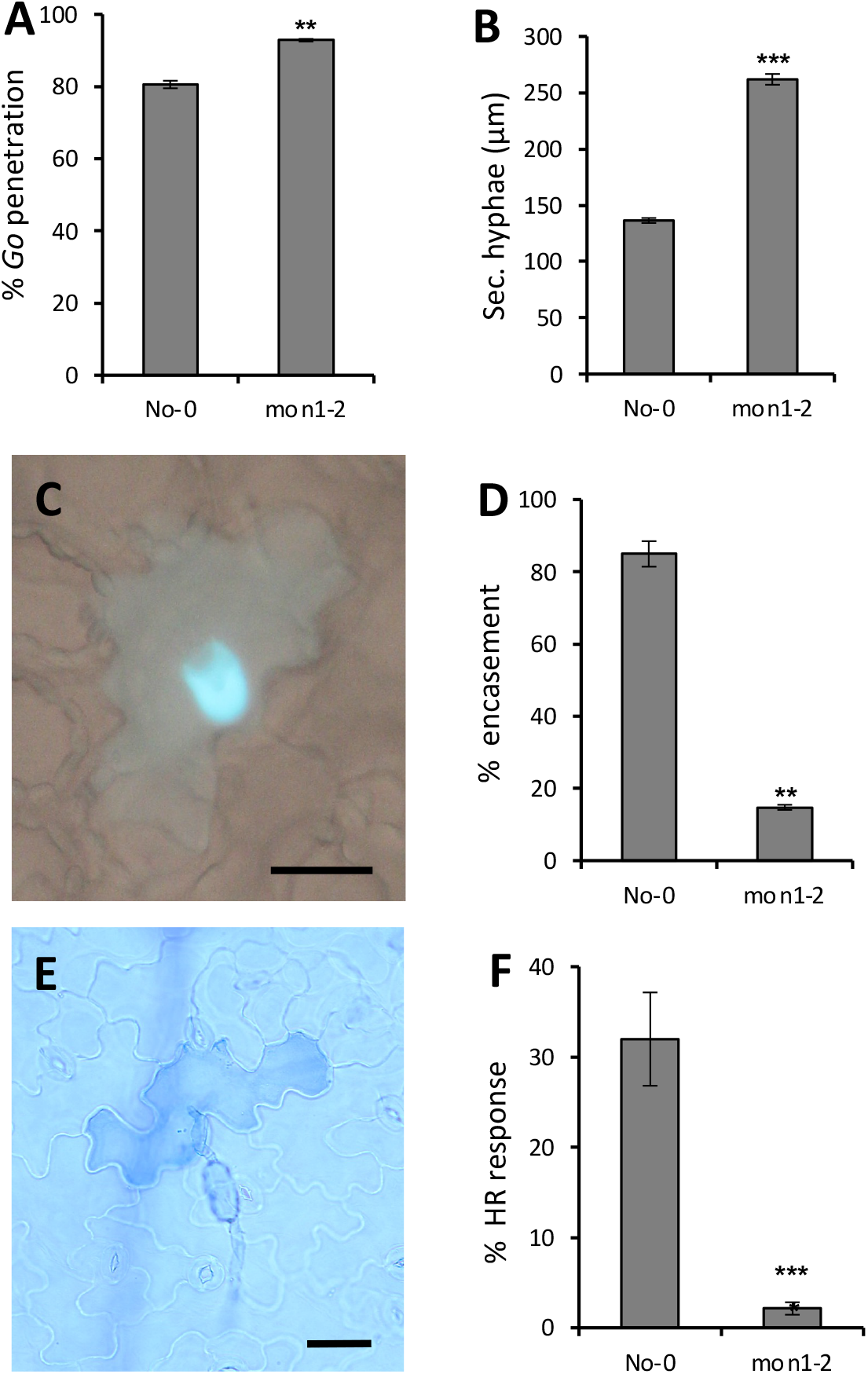
*At*MON1-dependent No-0 resistance *to Go* manifested as encasement formation and HR. **(A)** *mon1-2* caused marginally increased fungal penetration. **(B)** The *mon1-2* mutation allowed the fungus to develop longer secondary hyphae 2 days after inoculation. **(C-D)** The frequency of encasements around *Go* haustoria was strongly reduced in *mon1-2*. **(E-F)** HR of single epidermal cell attacked by *Go* was strongly reduced in *mon1-2*. All images and quantifications were made 2 days after inoculation. Size bars, 50 μm. Error bars, SE. **, *P*<0.01 and ***, *P*<0.001 assessed by Student’s T-tests. n=3.

### Hv*MON1 is required for barley Mla3-mediated resistance to* Bgh

Previously, disease resistance and HR mediated by the coiled-coil NLRs (CNLs), RPM1 and RPS2 were found to be dependent on the MVB-components AMSH3 and VPS4 in Arabidopsis (Schultz-Larsen *et al*., 2018). This and the finding that *Go* induces a MON1-dependent HR in No-0, inspired us to analyze whether *Hv*MON1 affects an Mla-mediated resistance to *Bgh* in barley, since these R-proteins are CNLs (Seeholzer *et al*., 2010). For this purpose, we employed virus-induced gene silencing (VIGS) of *Hv*MON1 in the barley line P-02, harboring the *Mla3* allele. The resulting plants had a 65% reduction in the *MON1* transcript levels (Fig. 5A) and were slightly smaller than the controls (Fig. 5B-C), but otherwise amenable for study. Interestingly, in *Hv*MON1 silenced plants the *Mla3*-mediated resistance was strongly suppressed, and the level of disease reached the level of that of susceptible plants without *Mla3* (Fig. 5D-E). Thus, *Mla3*-mediated resistance requires *Hv*MON1 and supports the importance of MVBs functioning in CNL-mediated resistance.

**Figure 5.**
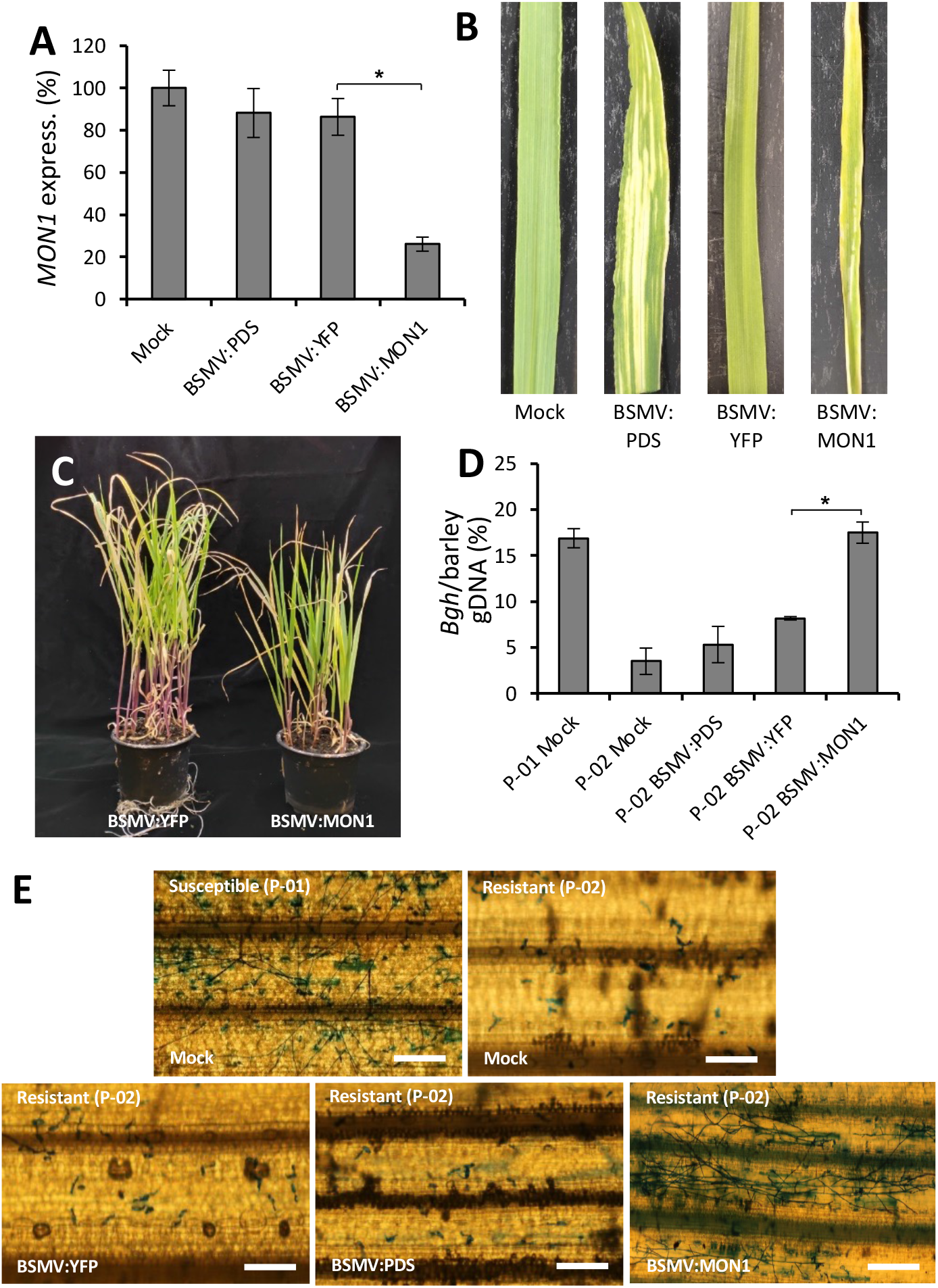
*Mla3*-mediated resistance to *Bgh* requires *Hv*MON1. **(A-C)** Barley stripe mosaic virus (BSMV)-induced gene silencing (VIGS)-based knock-down of the *HvMON1* transcript (A), reduces leaf width (B) and plant size (C). Samples taken from second leaves 14 days after virus inoculation of first leaves. Bleaching after VIGS-based knock-down of the phytoene desaturase (*PDS*) transcript and activation of *MON1* RNA-silencing using the YFP coding sequence serve as controls. (A) qPCR of the *HvMON1* transcript relative to the ubiquitin conjugating factor (UBC2) transcript. **(D-E)** The *Mla3*-mediated resistance in barley P-02 to the *Bgh* powdery mildew fungal isolate A6 is brokken by *HvMON1* knock-down. Four days after inoculation. Size bars, 100 μm. (D) *Bgh* qPCR-based quantification using *Bgh* glyceraldehyde-3-phosphate dehydrogenase and barley UBC2 genomic sequences. The data are mean values of three repeats. Error bars, SE. *, *P*<0.05 assessed by Student’s T-tests. n=3.

### *Germination defect and lethality of Col-0* mon1-1 *is partly immunity-dependent*

To counteract that effectors remove or inactivate immune components vital for defense, NLRs often monitor potential targets of effectors either directly or indirectly. Consequently, removal of guarded immune components by mutations may activate NLRs inappropriately and cause severe developmental phenotypes (Thordal-Christensen, 2020). Finding that *Hv*MON1 is an effector target of CSEP0162 and that MON1 plays a conserved role in plant immunity, we speculated if the rather severe lethality phenotype described for Col-0 *mon1-1*, could be a result of such a secondary activation of immunity (Cui *et al*., 2014; Ebine *et al*., 2014; Singh *et al*., 2014). In Arabidopsis, most sensor NLRs are of the TIR-NLR (TNL) type, the rest being CNLs (Meyers *et al*., 2003; Ngou *et al*., 2022). While TNL immune activation always depends on EDS1, CNL immune activation can depend on NDR1 (Aarts *et al*., 1998; Lapin *et al*., 2019). We therefore crossed the *eds1-2* and *ndr1-1* mutations into the Col-0 *MON1/mon1-1* heterozygous line. Next, we produced seeds of Col-0, Col-0 *MON1/mon1-1*, Col-0 *MON1/mon1-1 eds1-2* and Col-0 *MON1/mon1-1 ndr1-1* plants grown together under the same conditions and compared their germination rate (Fig. 6A). Seventy-seven point nine percent of the seeds of Col-0 *MON1/mon1-1* germinated, which is only marginally higher than the three-fourths expected if all homozygous *mon1-1* seeds failed to germinate. This result agrees well with previous findings (Singh *et al*., 2014). Moreover, the germination rate of Col-0 *MON1/mon1-1* seeds in the *ndr1-1* background was not different from that of wild type. Meanwhile, in the *eds1-2* background the Col-0 *MON1/mon1-1* seeds had a significantly higher germination rate (Fig. 6A). In addition, the improved germination was corroborated by the observation that *mon1-1 eds1-2* plants had longer roots and larger aerial parts than *mon1-1* plants (Fig. 6B-C). Overall, these observations show that the lethality inflicted by loss of MON1 is partly due to the activity of EDS1, which suggests that autoimmunity is activated by one or more TNLs.

**Figure 6.**
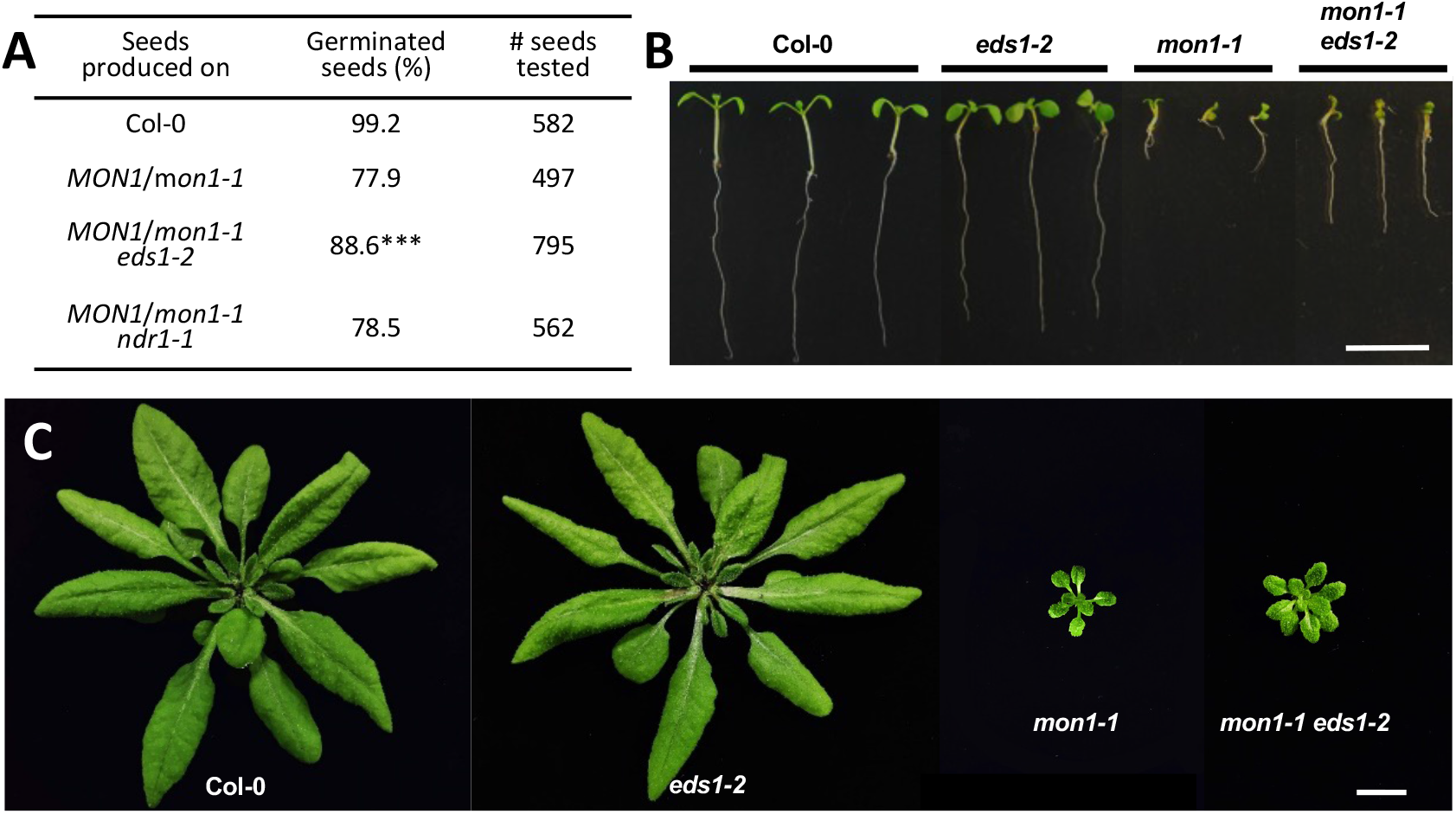
Lethality and petiteness of Col-0 *mon1-1* is partly EDS1-dependent. **(A)** Germination rates of seeds segregating for *mon1-1* in wild-type, *eds1-2* and *ndr1-1* backgrounds. **(B,C)** Root and rosette development affected by *mon1-1* and *eds1-2. mon1-1* and *mon1-1 eds1-2* plants were superior examples selected from (A), and their genotypes were confirmed by PCR. Plants in B are 10 days old (plate grown), and in C they are 5 weeks old. Size bars, 1 cm. *P*-values assessed by Chi-squared tests. ***, *P*<0.001.

## Discussion

In simple terms, papillae block penetration and encasements prevent exchange of compounds between the haustorium and the plant cytosol. In barley attacked by *Bgh*, encasement formation *per se* is not observed. Yet, early transmission electron microscopy (TEM) suggested that rudiments of encasements appear to be present (*e.g*. Heitefuss and Ebrahim-Nesbat, 1986). In this work, the authors refer to a “collar” present around the haustorial neck, which is very electron-light and distinct from the electron-dense papillae. In the meantime, they also describe some papillae that are two-layered – electron-dense in the first formed layer, and electron-light in the layer formed later. This electron-light layer is in other cases continuous with the collar. Nielsen et al. (2017) described that in *Bgh*-attacked Arabidopsis both papillae and encasements are labelled with PEN1, while only encasements are labelled with a constitutively active form of a Rab5 GTPase (ARA7^QL^). In the same study, encasement formation was shown to be ARA7-dependent, while Böhlenius *et al*. (2010) and Nielsen *et al*. (2012) showed that papilla formation is formed by a separate pathway, dependent on the ARF-GEF, GNOM, and the ARFA1b/c GTPases. These observations agree with the electron-dense structure in the TEM study of Heitefuss and Ebrahim-Nesbat (1986) being the papilla and the electron-light structure being the encasement. This in turn suggests that an encasement pathway indeed is activated in barley attacked by *Bgh*, but that it is somehow hampered. Furthermore, since this electron-light material is deposited as a layer on top of the papilla, it may contribute to blocking penetration by *Bgh*.

We can now add that MON1 is important for the encasement formation, and that the *Bgh* effector CSEP0162 interacts with MON1 and at the same time inhibits encasement formation in barley. Previously, Ahmed *et al*. (2015) obtained a 40% reduction in *Bgh*’s penetration rate by knock-down of CSEP0162 using host-induced gene silencing (HIGS). In this work, the effect of CSEP0162 on encasement formation was not considered. However, we speculate that the observed reduced penetration rate may have related to a fortification of the papilla because of less CSEP0162-mediated inhibition of the encasement pathway (Fig. 7). In Arabidopsis, encasements around *Bgh* and *Go* haustoria are labelled with the PEN1 syntaxin, known to associate with EVs (Nielsen *et al*., 2017; Rutter and Innes, 2017). As the secretion of PEN1 into the encasement matrix depend on activation of ARA7 by its GEF, VPS9a, it suggests involvement of MVBs fusing to the PM. Hereby, MVB intraluminal vesicles, labelled with PEN1, are secreted into the encasements as EVs, as suggested in the model in Fig. 7. Our present finding of MON1 being essential for encasement formation is very much in line with these previous findings, and it supports the idea that MVB fusion to the PM is required. We envisage that the MON1/CCZ1 complex is required to activate a RabG GTPase that may tether MVBs to the EXO70B2 complex as suggested by Ortmannová *et al*. (2022). We also found that MON1 is important for reducing the penetration rate, which also requires VPS9a and the EXO70B2 complex, as described by Nielsen *et al*. (2017) and Ortmannová *et al*. (2022). We believe this to be due to the encasement pathway’s contribution of another layer onto the papillae as indicated by TEM study of Heitefuss and Ebrahim-Nesbat (1986).

**Figure 7.**
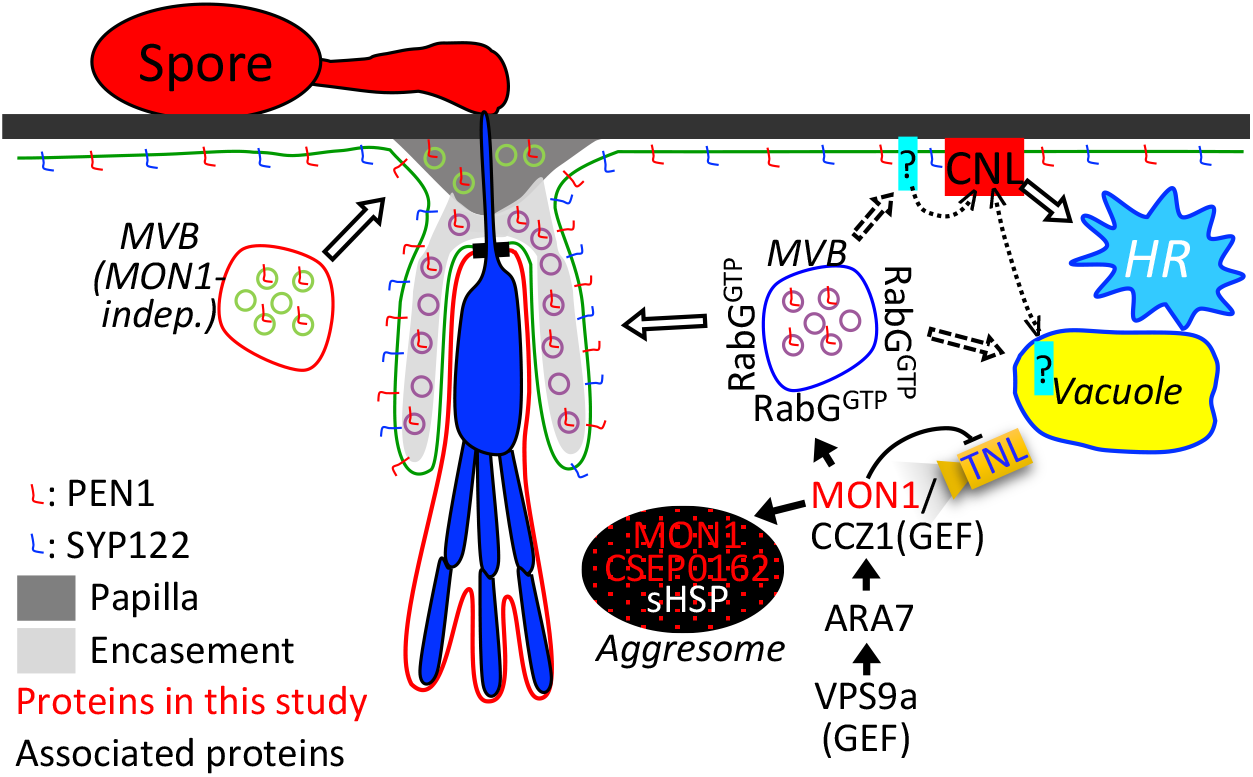
Model for the role of MON1 and *Bgh* effector CSEP0162 in immunity.

The MON1-dependence of Mla3-mediated HR described here, as well as previous findings of the MVB ESCRT components, AMSH3 and SKD1 (VPS4), being required for resistance and HR activated by the Arabidopsis CNLs, RPM1 and RPS2 (Schultz-Larsen *et al*., 2018), strongly suggests that MVBs serves a general function in CNL-activated HR. Therefore, according to the iceberg model (Thordal-Christensen, 2020) it would not be surprising if CSEP0162-inflicted inhibition of MVB fusion with the PM or tonoplast could prevent unknown CNLs from activating HR. Reversely, silencing CSEP0162 by HIGS may stimulate HR of *Bgh*-attacked cells and thereby reduce haustorium formation. As this was not addressed by Ahmed *et al*. (2015), future studies should test this hypothesis. Whether the MON1-requiring HR response in No-0, discovered here against *Go*, is activated by a CNL remains to be studied. In recent years, cell death activated by certain CNLs has been found to be caused by a homopentameric CNL complex, where the N-terminus of each CNL together forms a pore in the plant cell PM, through which a deleterious Ca^2+^ influx occurs (Wang *et al*., 2019; Bi *et al*., 2021). It will be interesting in the future to pursue whether MVB traffic to the PM plays a role in this process (Fig. 7), as it has not escaped our attention that both the Mla10 N-terminus (identical to the Mla3 N-terminus, Seeholzer *et al*., 2010) and the RPM1 N-terminus appear to be able to form such PM pores (Adachi *et al*., 2019). Alternatively, CNLs may depend on MVBs to send a negative regulator for degradation in the vacuole.

Arabidopsis membrane trafficking mutants often have growth defects due to loss of vital protein functions. Examples of this are *vps4* (*skd1*) (Haas *et al*., 2007), *vps9a* (Goh *et al*., 2007) and *gnom* (Mayer *et al*., 1991). Nonetheless, our observation that the *eds1-2* mutation partially rescues Col-0 *mon1-1* KO suggests that autoimmunity adds to the severity of the phenotypic growth defect. Similarly, *eds1-2* also rescues *amsh3* mutants and therefore links autoimmunity with loss of normal MVB function (Schultz-Larsen *et al*., 2018). It is becoming increasingly clear that immunity often is dependent on fundamental cellular processes such as membrane trafficking (e.g. Ortmannová *et al*., 2022; Rubiato *et al*., 2022), which therefore are potential effector targets. We also know that the NLR-based surveillance system monitors components targeted by pathogens and kills the cell when activated. Thus, we should expect membrane trafficking mutants to suffer from autoimmunity. Interestingly, the *mon1-2* KO in No-0 performs significantly better than Col-0 *mon1-1* KO (Cui *et al*., 2014). We speculate that this difference is due to the highly variable NLR populations present in the two ecotypes (Van de Weyer *et al*., 2019).

One arising question is how CSEP0162 may suppress MON1 function. This effector was previously found to interact with two sHSPs (Ahmed *et al*., 2015), and we found that CSEP0162 and MON1 co-localize in diffuse structures, not observed when these two proteins are expressed separately. We speculate if these structures are aggresomes similar to those formed when sHSPs interact with misfolded proteins (Johnston and Samant, 2021; Reinle *et al*., 2022). Thus, we suggest that CSEP0162 links MON1 to sHSPs that subsequently induce formation of aggresomes containing all three proteins (Fig. 7). In fact, at an early stage of the interaction, sHSPs may attract larger HSPs to CSEP0162 and MON1, which targets them for ubiquitination and proteasomal degradation, or aggresomes may alternatively be removed by autophagy (Johnston and Samant, 2021; Reinle *et al*., 2022), thereby preventing MON1 function. A twist here is that MON1 itself is required for autophagy (Hegedüs *et al*., 2016) suggesting that removal of the aggresome in this case may be difficult.

In conclusion, our data show that MON1 plays important roles in encasement formation as well as in HR responses and that the *Bgh* effector CSEP0162 interacts with this protein important for immunity. We suggest that CSEP0162 by this interaction and its additional interaction with sHSPs diverts MON1 into aggresomes and potential degradation. With such Swiss army knife-like properties, we believe that MON1 and CSEP0162 are central to barley’s interaction with the powdery mildew fungus, and that this effector is an important reason why we see little encasements and HR in compatible barley-*Bgh* interactions.

## Acknowledgements

We thank Professor Gerd Jürgens for *mon1-1* and Professor Jane Parker for *eds1-2*. Associate Professor Shoi Mano for the BiFC-vectors and Professor Dawei Li for providing the BSMV-RNAi system. Research support has been obtained from China Scholarship Council (PhD-student scholarship 201706850092 to WL), Novo Nordisk Foundation (Challenge grant NNF19OC0056457 to HTC), and Villum Fonden (Experiment Programme grant 00028131 to HTC).

## Author contributions

Wenlin Liao: Performed the work and generated many concepts

Mads E. Nielsen: Supervised the work

Carsten Pedersen: Supervised the work

Wenjun Xie: Supervised the confocal imaging and genetics

Hans Thordal-Christensen: Supervised the work and wrote the first manuscript version

---

**Supplementary Figure S1. Callose-containing encasements around *Bgh* haustoria in barley induced by tetraconazole.**

**Supplementary Figure S2. Barley *Hv*MON1 complements the function of Arabidopsis *At*MON1.**

**Supplementary Video S1. Co-expression of mYFP-CSEP0162 and mCherry-*Hv*MON1 in barley P-02 protoplasts observed by LSCM 20 h after transformation.**

**Supplementary Table S1. Primers used in this work.**

**Supplementary Table S1. Gateway destination vectors used in this work.**

